# MethFlow^VM^: a virtual machine for the integral analysis of bisulfite sequencing data

**DOI:** 10.1101/066795

**Authors:** Ricardo Lebrón, Guillermo Barturen, Cristina Gómez-Martín, José L. Oliver, Michael Hackenberg

## Abstract

The analysis of whole genome DNA methylation patterns is an important first step towards the understanding on how DNA methylation is involved in the regulation of gene expression and genome stability.

Previously, we published MethylExtract, a program for DNA methylation profiling and genotyping from the same sample. Over the last years we developed it further into a methylation analysis pipeline that allows to take full advantage of novel genome assembly models. The result is a new pipeline termed MethFlow which permits both, profiling of methylation levels and differential methylation analysis.

Frequently DNA methylation research is carried out in the biomedical field, where privacy issues play an important role. Therefore we implemented the pipeline into a virtual machine termed MethFlow^VM^ which shares with a web-server its user-friendliness however, the decisive advantage is that the sequencing data does not leave the user desktop or server and therefore no privacy issues do exist.

The virtual machine is available at: http://bioinfo2.uqr.es:8080/MethFlow/

## Introduction

Next generation sequencing techniques triggered substantial changes in many research fields of molecular biology. Specific protocols allow now the fast and cheap expression profiling of whole transcriptomes, the detection of transcription factor binding sites, histone modifications and whole genome methylation maps among many other analysis types [1]. Especially research on DNA methylation was boosted as before whole genome studies were hampered by the fact the PCR does not copy the methyl group and hybridization is insensitive to the methylation state of the cytosine [2], DNA methylation is an essential epigenetic mark involved in the regulation of gene expression and genome stability [3–5]. The analysis of whole genome DNA methylation patterns has become an important first step towards the understanding on how DNA methylation exerts its functions in the development of an organism and why dysregulation is frequently linked to pathologies like cancer [6].

Currently, most high-throughput sequencing protocols for DNA methylation profiling rely on the pre-treatment of the denaturalized DNA with sodium bisulfite which coverts un-methylated Cytosine first into Uracil and finally into Thymine. After sequencing and alignment against a reference genome, the methylation states can be read out of the read alignments. Many different algorithms have been developed for this task like BSseeker [7] Bismark [8], NGSmethPipe [9] or MethylCoder [10]. Furthermore, some of the available programs allow additionally the detection of sequence variants from the same sample like Bis-SNP [11] or Methyl Extract [12].

Many DNA methylation research is currently carried out in biomedical and clinical research fields. This implies that frequently the sequencing data underlies privacy issues what rules out the usage of web-servers and probably even web-based platforms like Galaxy [13]. As a consequence, the data needs to be analysed locally what requires some infrastructure and system administrators. The asset costs and administration of Linux machines together with the need for personal trained in HTS-data analysis raises the cost for HTS-based research enormously bringing especially smaller groups or laboratories into difficulties. One possible solution are preconfigured virtual machines, which can be run on any operating system, and no system administration and maintenance is needed.

We present MethFlow^VM^, a virtual machine for DNA methylation profiling, genotyping and differential methylation analysis. The virtual machine is based on an extended version of our MethylExtract algorithm [12], Bismark [8] /Bowtie2 [14] for read alignment and methylKit [15] for differential methylation analysis. The virtual machine can be easily used by users without bioinformatics background.

## Scope of MethFlow^VM^

Virtual machines can be executed on all common operating systems through virtualization software like VirtualBox (https://www.virtualbox.org/), VMware (http://www.vmware.com/), Vagrant (https://www.vagrantup.com/) or XenServer (http://xenserver.org/). MethFlow^VM^ is the implementation into a virtual machine of two basic analysis types: i) profiling of cytosine methylation values through the analysis of bisulfite sequencing data and ii) detection of differential methylation either at a cytosine or region level. The download size of the virtual machine is currently 1.6 Gb (without desktop) and we recommend at least 4 CPUs and 8 Gb free memory for the host computer. A 64-bit processor is required. In the tutorial we explain in detail how the virtual machine can be obtained and started. MethFlow^VM^ implements several small tools that help the user to automatically set up, maintain and update the virtual machine.

## DNA methylation workflow

The workflow of DNA methylation profiling can be seen in Figure 1. The data analysis can be started at different levels depending on the kind of input data. Briefly, the pipeline performs the following steps:

- **Format conversion**: Conversion of SRA data format into FASTQ. by means of the SRA Toolkit. This only applies if the input data comes from the Sequence Read Archive (SRA) public repository [16] which is in ‘sra’ format and needs to be converted first.
- **Adapter Trimming and low quality reads**: if the input data is in fastq format, the user can either chose to trim the adapter sequences or to omit this step. The adapter trimming is performed by means of Trimmomatic [17]. Furthermore, Trimmomatic is used to remove low quality 3’ ends with ‘trailing’ mode applying by default a PhredScore threshold of 28 (Q <= 28). Reads shorter than half of the original read length are eliminated (MINLEN option). All parameters can be changed by the user.
- **Alignment to the genome**: Adapter trimmed input reads in fastq format (either provided by the user or generated in the previous step) are aligned to a three letter genome by means of Bismark [8] which uses Bowtie2 [14] as aligner. The output of this step is a standard BAM file.
- **Elimination of known technical artefacts**: different technical artefacts can bias the methylation profiling results like overhang end-repair, 5’- bisulfite conversion failure, undetected adapter sequences and sequencing errors. BSeQC [18] detects automatically regions at the 5’ and 3’ end whose DNA methylation values might be biased by any technical artefact. BSeQC trims each read at the 5’ end by a given number of nucleotides – those putatively affected by technical artefacts. In ‘normal’ samples, not more than 10 nt should be trimmed of.
- **Detection of DNA methylation and genotype**: the DNA methylation profiling and genotyping is performed by means of MethylExtract [12]. The program takes into account several important error sources like sequencing errors, bisulfite failure, clonal reads, and single nucleotide variants in order to generate high quality whole genome methylation maps.
- **Graphical representation**: The obtained results can be visualized in several ways. For basic summary graphics like the distribution of the methylation values or read coverage, MethFlow^VM^ implements methylKit [15]. The workflow converts automatically MethylExtract format into methylKit format and generates the mentioned plots. Note that the automatic conversion of MethylExtract into methylKit format is the first step into the differential methylation analysis which can be performed as well with methylKit.
- **Downstream analysis**: MethFlow^VM^ provides several useful auxiliary programs for downstream analysis. First, the cystosine methylation data can be easily visualized by means of the UCSC genome browser[19] or IGV [20]. MethFlow implements the required programs that convert MethyExtract format into BedGraph, BED6, BED6+6, bigBed or bigwig. Second, MethFlow^VM^ provides a program for the automatic upload of methylation results to Galaxy [21] which can then be used for further downstream analysis. Note that those programs are not part of the methylation workflow as they are not launched automatically. We explain in the manual step by step how those programs can be used.

**Figure 1:**
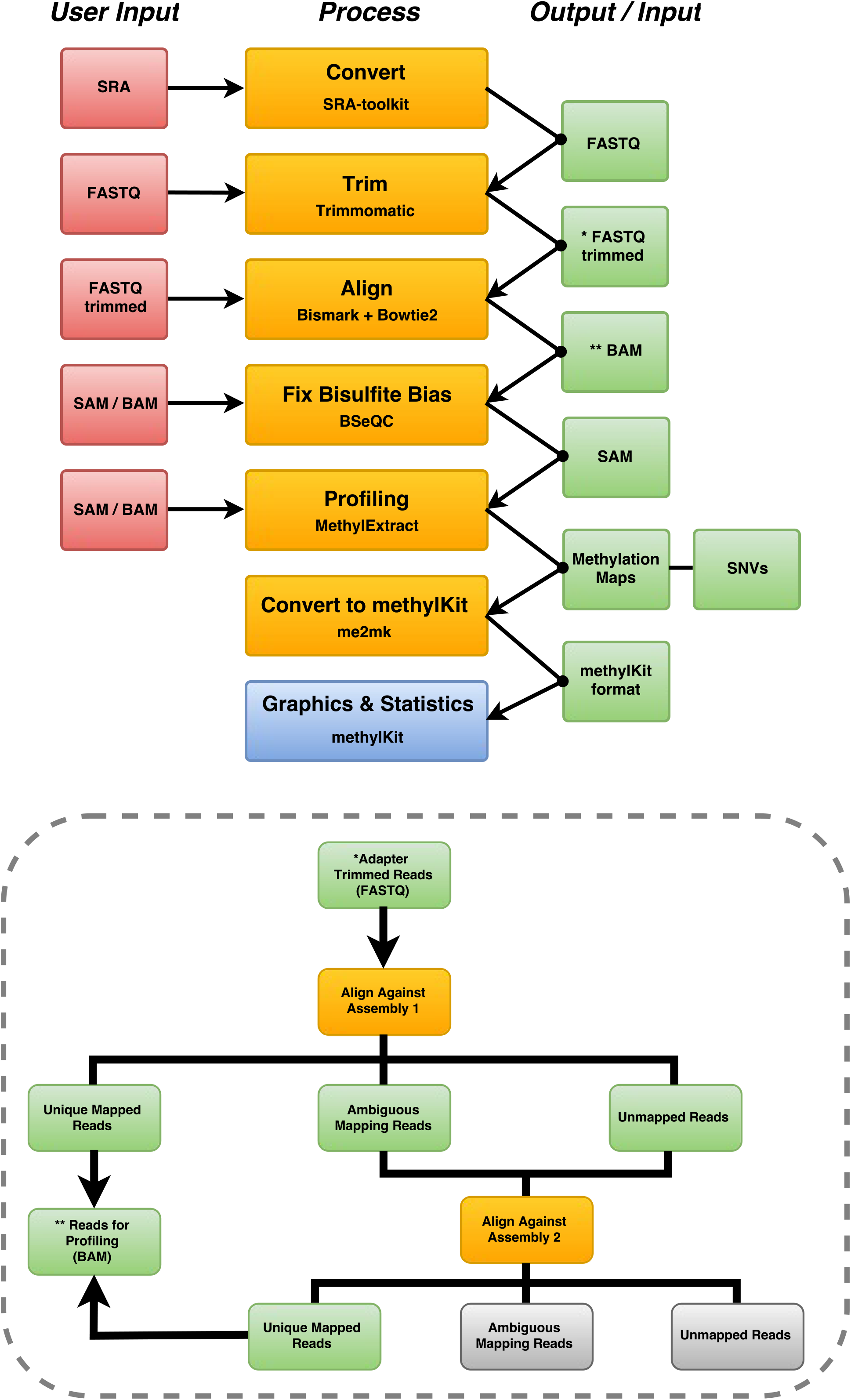
Workflow of the methylation pipeline. On the left side, the accepted user input formats are depicted and on the right side the intermediate formats produced by the different components in the workflow. Depending on the input data, the workflow starts at a different level. The last step of the pipeline consists in the conversion of MethylExtract format into methylKit format which can be directly used for differential methylation analysis. The inlay on the bottom depicts the novel two-step mapping strategy against two different assemblies. As shown, in the second mapping the user can chose either to map ambiguous read, unmapped reads or both to the second assembly. For example, the first assembly could be the reference assembly plus all decoy sequences while the second assembly represents only the canonical reference sequences (see ‘New features in MethFlow’).

## New features in MethFlow

The reference assembly is crucial for the analysis of sequencing data in general. However, constructing a reference assembly by collapsing sequences from several individuals into a single consensus haplotype representation of each chromosome as done normally has several drawbacks [22]. To overcome the limitations of such a sequence model, the GRC (Genome Reference Consortium) started with the inclusion of ‘alternative sequence paths’ in the GRCh37 (hg19) assembly [23]. Those new ‘sequence paths’ are introduced to reflect the existing sequence and structural variation within a given population. It can be expected that the inclusion of these alternative sequences increases the number of correctly mapped reads. One important consequence of those new assembly models is that they are neither haploid nor diploid, but they contain additional sequences for regions with a high degree of diversity in the population. Normally programs expect a haploid representation and therefore those new models pose additional challenges to the programs used for the analysis of sequencing data. Especially in DNA methylation profiling those issues are important as only uniquely mapping reads are used and multiple-mapping reads are discarded. Furthermore, incorrectly mapped reads will lead to incorrect methylation profiling and sequence variation between the reference sequence and the sample haplotype, which pose additional challenges in bisulfite sequencing analysis. Therefore, especially in the analysis of WGBS data analysis the inclusion of alternative sequences will improve the mapping quality and therefore the quality of the methylation maps. However, when using only one assembly that incorporates those additional sequences (decoy sequences), very likely many reads will map multiple times within the variable regions - and those reads are lost for the analysis.

In order to i) exploit the advantages of the new assembly models and ii) recover the useful information of multiple-mapped reads for the analysis, we incorporated a twostep mapping process. First, the reads are mapped against a decoy assembly (canonical chromosomes + decoy sequences). This increases the number of correctly mapped reads but also the number of multiple-mapped reads. A certain percentage of those ambiguous reads can be recovered in a second mapping step against the canonical chromosomes (see bottom inlay of Figure 1).

Note that this two-step mapping procedure can be applied in different scenarios. For example, if a population specific assembly exists whose haplotype structure is closer to the analysed sample, but of lower quality than the reference assembly. In such cases the population specific assembly can be used in the first step and the reference assembly in the second.

## Protocol and working example

To show the usefulness of MethFlow^VM^ we generated 9 small test datasets which can be downloaded easily from within the virtual machine. The manual http://bioinfo2.ugr.es:8080/MethFlow/manual/ contains a step-by-step protocol on how to obtain the data and how to analyse it by means of the virtual machine. The test data set was obtained from the ROADMAP Epigenomics Project [24]. In concrete we used data from three different tissues (adipose, psoas muscle and small intestine) and from three different individuals (STL001, STL002 and STL003). From these datasets we extracted the reads for two genome regions: i) chr12:7,600,000-7,700,000 (APOBEC1) and chr19:2,200,000-2,300,000 (AMH).

MethFlow can be launched in three different ways. The easiest is an interactive way by means of dialogs (see figure 2a), but it accepts also a direct input of parameters in the command line or through a configuration file. Furthermore, several arguments like the ‘base output directory’ or the indexes of the genome assemblies can be provided in a setting file (.methflowrc) and therefore they don’t need to be specified on every run. MethFlow^VM^ helps the user through the configuration when the VM is used for the first time. Especially we want to stress the importance of using shared folders between the host and the virtual machine which accelerates some steps of the analysis up to 4 times. All recommended configuration steps are explained step-by-step in the manual.

**Figure 2:**
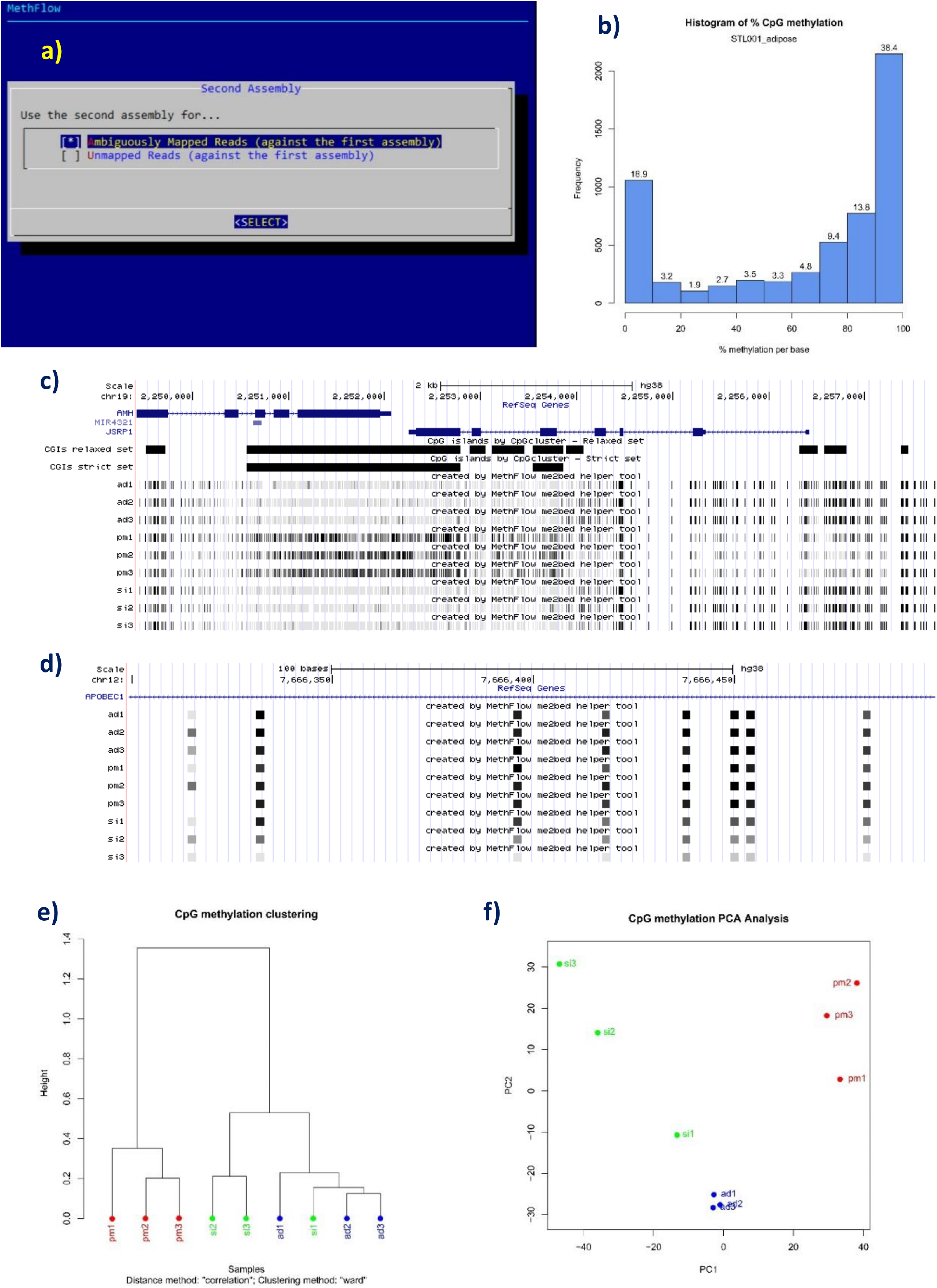
a) Dialog window guiding the user through the required input data. In this case, the user is asked if only multiple-mapped reads should be used for the alignment against the second assembly or if multiple and unmapped reads should be used instead, b) The distribution of methylation levels in adipose of the STL001 individual, c) The methylation levels of all CpGs within a region containing the AMH (Anti-Müllerian hormone) and JSRP1 genes. Apart from the visualization of the genes, the methylation level of each CpG within this region is shown (9 tracks at the bottom). Unmethylated CpGs are depicted by grey-shaded bars while methylated CpGs are painted in black. The differentially methylated CpG island overlapping both 3’UTRs can clearly be distinguished, d) Differential methylation of single CpGs can be seen in the small intestine (especially in STL003) in an intronic region of the gene APOBEC1. e) Cluster tree and f) PCA analysis generated by means of methylKit showing the tissue specificity of methylation patterns.

By typing ‘MethFlow run --use_assembly2’ in a terminal, a dialog pops-up which guides the user through the selection of the mandatory parameters. The ‘-use_assembly2’ parameter will launch the workflow with two assemblies (see section ‘New features in MethFlow’). Depending on the input data, up to 14 output folders are generated by the different components of the pipeline. Furthermore, MethFlow writes a file with the references of all used tools. Depending on the input data format, MethFlow launches sequentially a certain number of programs. The CITE.txt file contains the references to be cited if the results are used in a scientific publication.

Figure 2b-f show several results of MethFlow. Figure 2b shows the distribution of methylation levels of CpG cytosines in adipose of the STL001 individual. The typical U shaped distribution can be seen with 18.9% of CpGs having methylation levels lower than 0.1 (unmethylated CpGs) and 38.4% of CpGs show methylation levels higher than 0.9 (methylated CpGs). Figure 2c shows a sub-region of chr19:2,249,421-2,257,733 including the gene AMH (Anti-Müllerian hormone) and JSRP1, a tissue specific muscle gene involved in excitation-contraction coupling at the sarcoplasmic reticulum. Both genes are located ‘tail-to-tail’ having a very prominent CpG island overlapping the 3’ UTR’s of both genes. Interestingly, the CpG island is completely unmethylated (grey shaded bars) in adipose (ad) and small intestine (si) in all three individuals but strongly methylated (black bars) in psoas muscle (pm) in all three individuals. This might indicate that the methylation pattern of the muscle specific JSRP1 gene is involved in the regulation of the expression of this gene in an atypical way as the canonical regulation pathways are expected to act in the promoter region. CpG islands overlapping the 3’ end of genes were not extensively investigated so far but seemingly they are associated to cancer development [25]. Furthermore they are and characterized by an increased number of sites recognized by the Sp1 protein [26] and might therefore indeed play roles in the regulation of the corresponding genes.

Figure 2d shows part of an intron of the gene APOBEC1 (from the genomic region chr12:7,666,300-7,666,500). This gene encodes the protein Apolipoprotein B mRNA editing enzyme, catalytic polypeptide 1 also known as C->U-editing enzyme APOBEC-1. This enzyme is essential for the intestine to produce APOB48 instead of APOB100. APOB48 is produced exclusively in the intestine and is responsible for transporting chylomicrons. APOB100 is an apolipoprotein transporting cholesterol. The dysfunctionality of APOBEC1 can cause the development of hypercholesterolemia and atherosclerosis. Figure 2d shows a different methylation pattern of single CpGs between small intestine (si) and the other two tissues (ad: adipose; pm: psoas muscle), being less methylated in the former.

Finally we used the methylKit formatted methylation output files generated by MethFlow to perform a cluster analysis. Figure 2e shows the clustering tree and figure 2f the principal component analysis of the 9 samples. Both analyses confirm the tissue-specificity of the DNA methylation patterns as the samples are grouped by tissue type and not by genotype. MethylKit can be used also for a full differential methylation analysis, including single cytosines and regions. In the manual we discuss how the MethFlow output can be used directly as methylKit input to carry out those analysis.

## Conclusions and Future work

We present here a virtual machine, MethFlow^VM^ for the analysis of bisulfite sequencing data, including DNA methylation profiling, SNV calling and differential methylation analysis. A virtual machine shares with a web-server its user-friendliness as they can be used on any operating system without the need for system administration or maintenance. However, the decisive advantage is that the sequencing data does not leave the user desktop or server and therefore no privacy issues do exist.

Several new features are planned for the future or are already under development. Currently we are finishing a new version of our single cytosine resolution methylation database NGSmethDB [27, 28] which will include a collection of pairwise differentially methylation cytosines (DMC). The DMCs are calculated as the consensus of MOABS [29] methods and methylKit [15] and the incorporation of the consensus differential methylation method is scheduled for the next update of MethFlow^VM^. Furthermore we are already working in the full integration of NGSmethDB into MethFlow. The goal is to allow the user to profile the bisulfite sequencing data locally and compare it to all tissues and/or pathophysiological conditions stored in NGSmethDB without the necessity to upload the user data to any public servers. It will be possible to carry out the comparison by means of several programs providing differentially methylation at single cytosine and regional level. Specially for the differential methylation at a regional level we will incorporate several improvements like a novel segmentation algorithm adapted to the special requirement of DNA methylation derived from an existing method [30] and the detection of differentially methylation clusters applying WordCluster [31] and GenomeCluster [32] methods to differentially methylated cytosines.

## Acknowledgment

The research grant of the Ministerio de Economía y Competitividad from the Spanish Government [AGL2013-49090-C2-2-R to JLO and MH] is acknowledged.

